## Supplementary material for "C/EBPB-dependent Adaptation to Palmitic Acid Promotes Tumor Formation in Hormone Receptor Negative Breast Cancer": Supp Figures

**SUPPLEMENTAL INFORMATION**

**Supplementary Figure 1.**

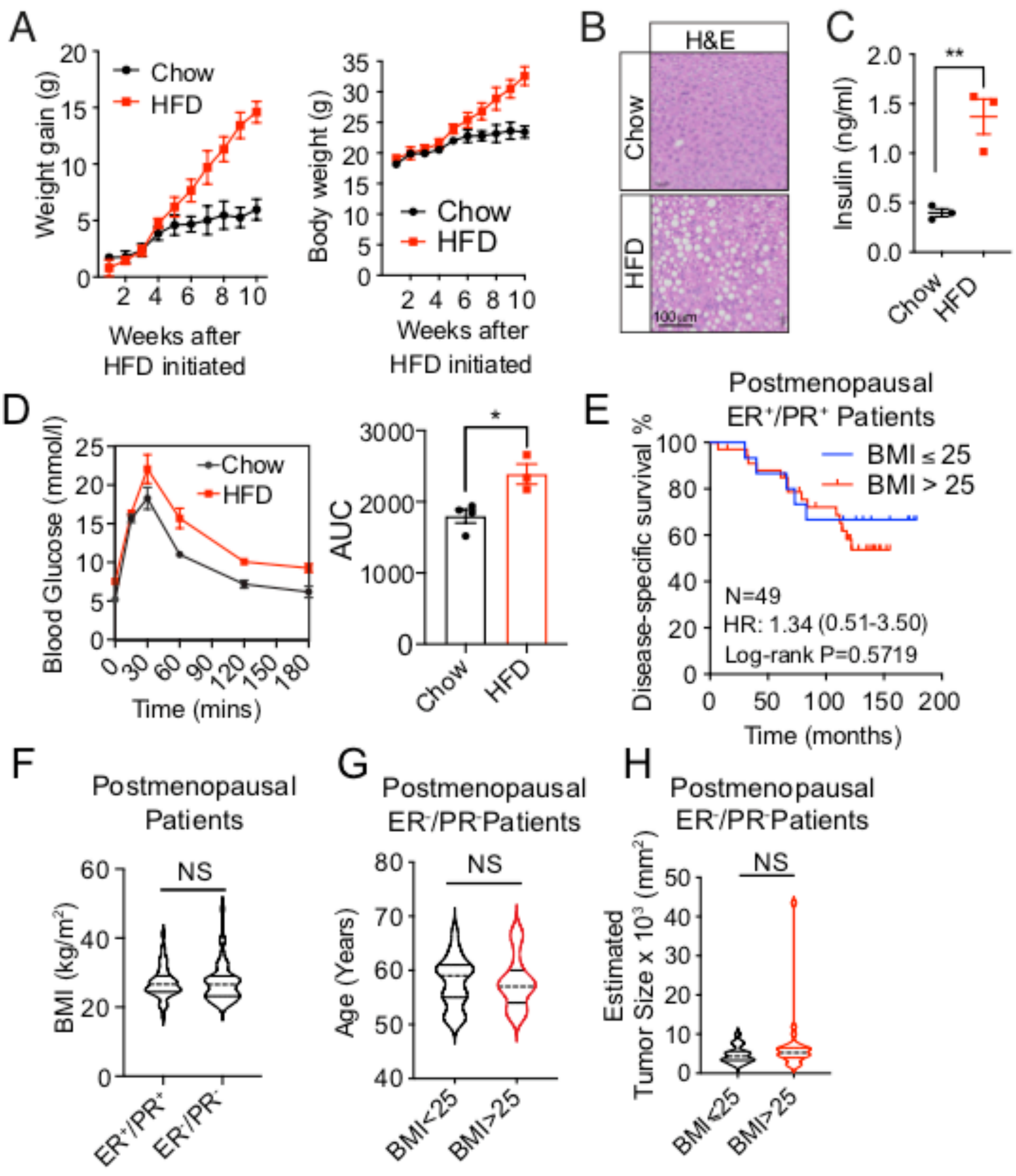


(A) Body weight gain (left panel) and absolute body weight (right panel) of HFD and chow-fed mice before implantation of tumors. Six-weeks old female C57BL/6J mice were started on HFD or standard chow diet (n=4 per group) for ten weeks prior to the tumor implantation. The measurement of animal body weight was started at six weeks of age and recorded weekly. For each time point, data is represented as mean ± SEM of four mice per group (n=4 per group).

(B) Hematoxillin and Eosin (H&E) stained tissue sections of livers from HFD and chow-fed mice. After ten weeks of HFD or chow diet feeding, female C57BL/6J mouse were sacrificed and and livers were harvested. Liver sections were stained using H&E. Histological analysis showed increased liver steatosis in mice from the HFD group compared to mice from the chow group.

(C) Concentration of fasting plasma insulin in HFD and chow-fed mice. Concentrations were determined by ELISA using overnight fasted blood samples collected from female C57BL/6J mice fed an HFD or chow diet for ten weeks (n=3 per group).

(D) Oral glucose tolerance test performed on HFD and chow-fed mice. Blood glucose clearance was determined in mice fed an HFD (n=3) or chow (n=4) diet for ten weeks. Blood glucose concentrations were measured at 0min, 15mins, 30mins, 60mins, 120mins and 180mins following glucose administration by oral gavage. For each time point, data is represented as mean ± SEM. AUC = area under the curve.

(E) Kaplan-Meier curves display disease specific survival for postmenopausal and $ER^+/PR^+$ patients (N=49) with high (red, BMI > 25) or low (blue, BMI ≤ 25) BMI. Log-rank (Mantel-Cox) P value is denoted for difference in disease specific survival. The analysis showed no significant difference between the groups.

(F) Distribution of BMI in postmenopausal ER+/PR+ and ER-/PR- patients. BMI distribution was similar between the groups.

(G-H) Distribution of postmenopausal $ER^-/PR^-$ patients' age (G) and estimated tumor size (H) in high (BMI > 25) and low (BMI ≤ 25) BMI groups. The estimated tumor size was calculated by multiplying the largest diameter by its perpendicular. The analysis showed no significant difference between the groups.

For C-D, statistical significance determined with unpaired, two-tailed Student's t-test. For F-H Kolmogorov-Smirnov test was used for statistical testing. (NS, P value > 0.05; *, P value < 0.05; **, P value < 0.01).

**Supplementary Figure 2.**

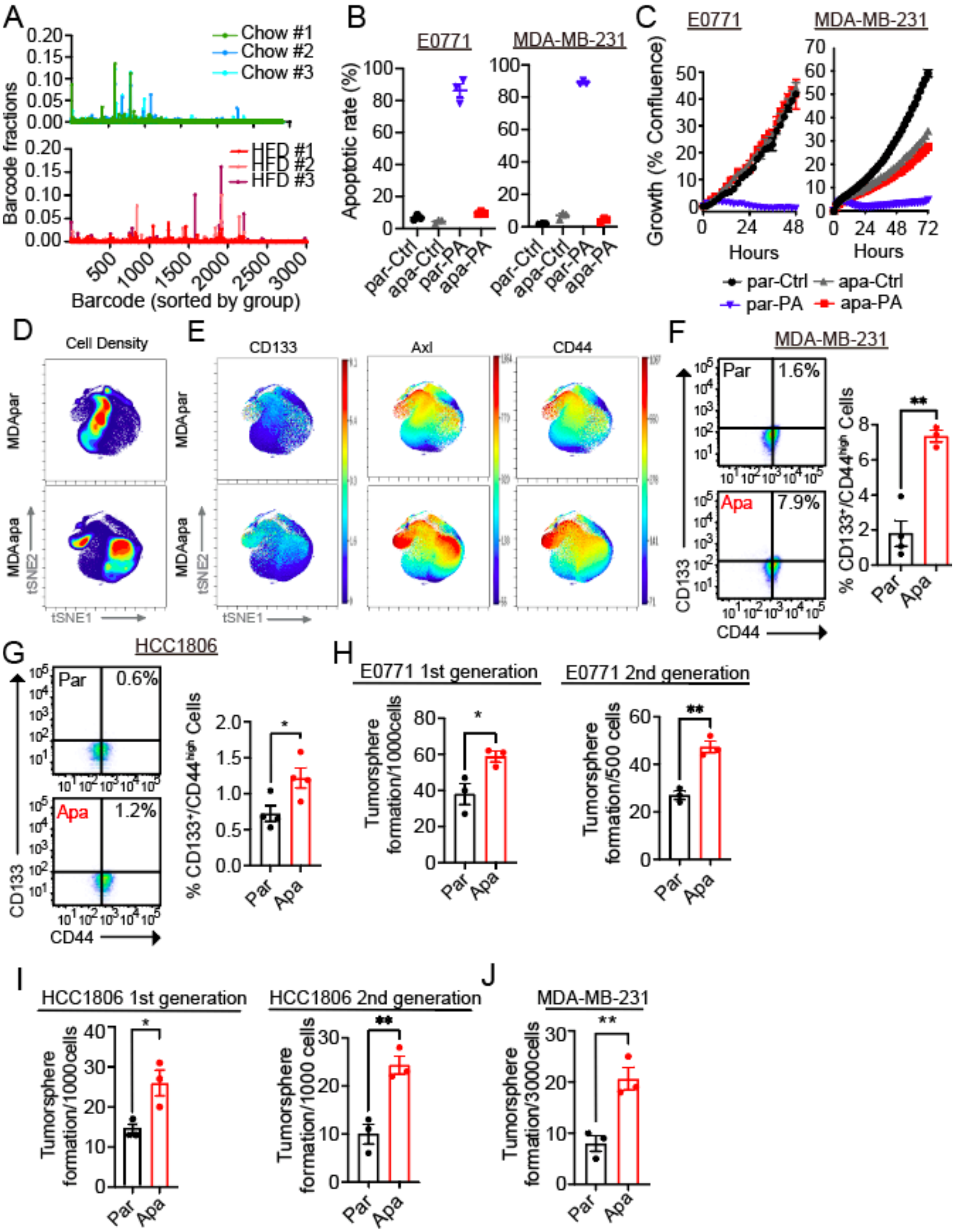

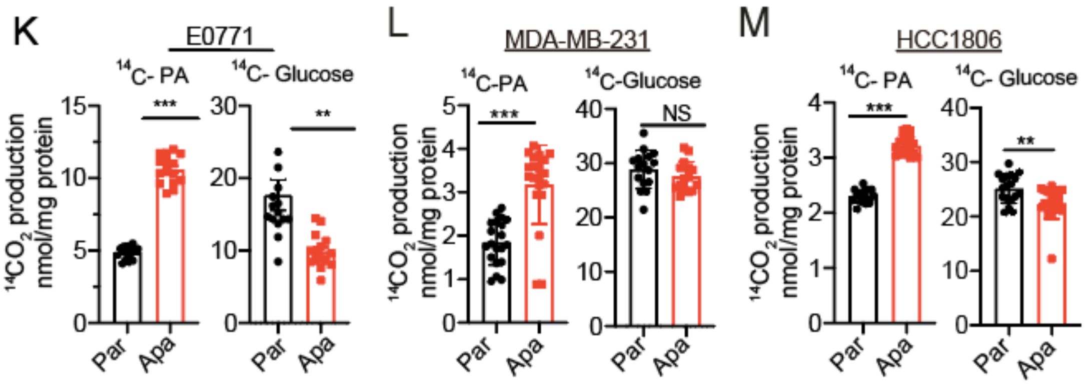


(A) Barcode distribution of all replicates of tumors derived from chow (upper panel) and HFD (lower panel) mice. The x axis of the histograms is barcode ID which were sorted by group and each bar represents one unique barcode.

(B) Apoptotic rate of parental and adapted E0771 (left panel) and MDA-MB-231 (right panel) cells that were treated with PA (500 μM for E0771 and 400 μM for MDA-MB-231) and vehicle (Ctrl) for 48hrs. Data are represented as mean ± SEM of three replicates.

(C) Time-dependent proliferation assay of parental and adapted E0771 (left panel) and MDA-MB-231 (right panel) cells following 48-72hrs. Cells were exposed to 400 μM (for MDA-MB-231 cells) or 500 μM (for E0771 cells) PA and vehicle (Ctrl). Cell growth was determined by high content imaging and represented as % confluence normalized to t=0. For each time point, data are represented as mean ± SEM of four to eight replicates.

(D) Representative contour plots of mass cytometry data colored by density of cells showing the changes between parental and adapted MDA-MB-231 cells. Total number of analyzed cells per cell line is equal to 100 000 cells. Color code represents the cell density from low (blue) to high (red).

(E) Representative tSNE plots of single parental and adapted MDA-MB-231 cells colored by expression of CD133, Axl and CD44.

(F-G) $CD133^{+}/CD44^{high}$ cells population in parental and adapted MDA-MB-231 (F) and HCC1806 (G) cells. Cells were stained by CD133-APC and CD44-FITC antibodies and measured by flow cytometry. Quantification data is shown as mean ± SEM of four replicates (one outlier in MDA-MB-231apa group was excluded from the quantification by Grubbs' outlier test).

(H-J) Tumorsphere formation assay and serial tumorsphere propagation assay of parental and adapted E0771 (H), HCC1806 (I) and MDA-MB-231 (J) cells. Cells (1000 cells/well for E0771 and HCC1806 cell lines, 3000 cells/well for MDA-MB-231 cell lines) were seeded into ultra-low attachment 6-well plates in the stem cell media and following 5-10days of growth, tumorspheres were imaged and quantified. The serial tumorsphere propagations were performed by dissociating the primary tumrospheres, and following the same method to reseed 500 cells/well for E0771 cells (H right panel) and 1000 cells/well for HCC1806 cells (I right panel). Quantifications of tumorspheres are represented as mean ± SEM of three replicates for each condition (n=3 / condition).

(K-M) Comparison of fatty acid and glucose oxidation assays between parental and adapted cells. Fatty acid oxidation on parental and adapted E0771 (K), MDA-MB-231 (L) and HCC1806 (M) cells was measured by cumulative $^{14}CO_2$-production during incubation with radio-labeled [1-$^{14}$C] palmitic acid (left panel). Glucose oxidation was shown by cumulative $^{14}CO_2$-production during incubation with radio-labeled D-[$^{14}$C(U)] glucose (right panel).

For F-M, statistical significance determined with unpaired, two-tailed Student's t-test. (NS, P value > 0.05; *, P value < 0.05; **, P value < 0.01; ***, P value < 0.001).

**Supplementary Figure 3.**

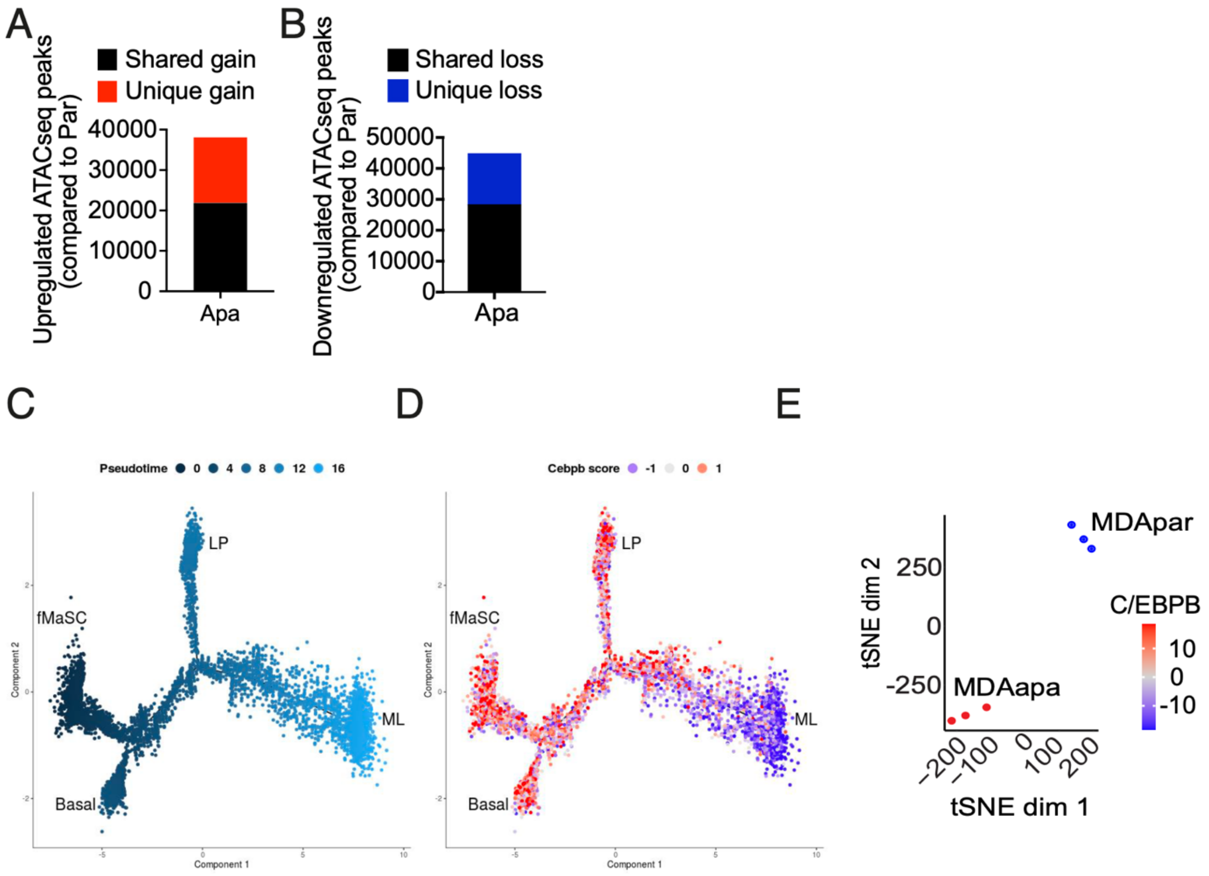


(A) Total number of significantly upregulated ATACseq peaks in MDAapa relative to MDApar with a FDR < 0.05. Unique gain peaks refer to peaks identified only in the adapted condition, whereas shared peaks are peaks called in both conditions.

(B) Total number of significantly downregulated ATACseq peaks in MDAapa relative to MDApar with a FDR < 0.05. Unique loss peaks refer to peaks identified only in the parental condition, whereas shared peaks are peaks called in both conditions.

(C) Pseudotime analysis of single-nuclei ATACseq of murine mammary cells at different developmental stages (GSE125523).

(D) Motif enrichment of transcription factors C/ebpb in the open chromatin regions at each individual cell along the mammary gland developmental trajectory is shown. fMaSC, fetal mammary stem cells; basal, adult basal cells; LP, luminal progenitors; and ML, mature luminal cells.

(E) t-SNE clustering of individual MDApar and MDAapa replicates showing differential motif enrichment in transcription factors C/EBPB.

**Supplementary Figure 4.**

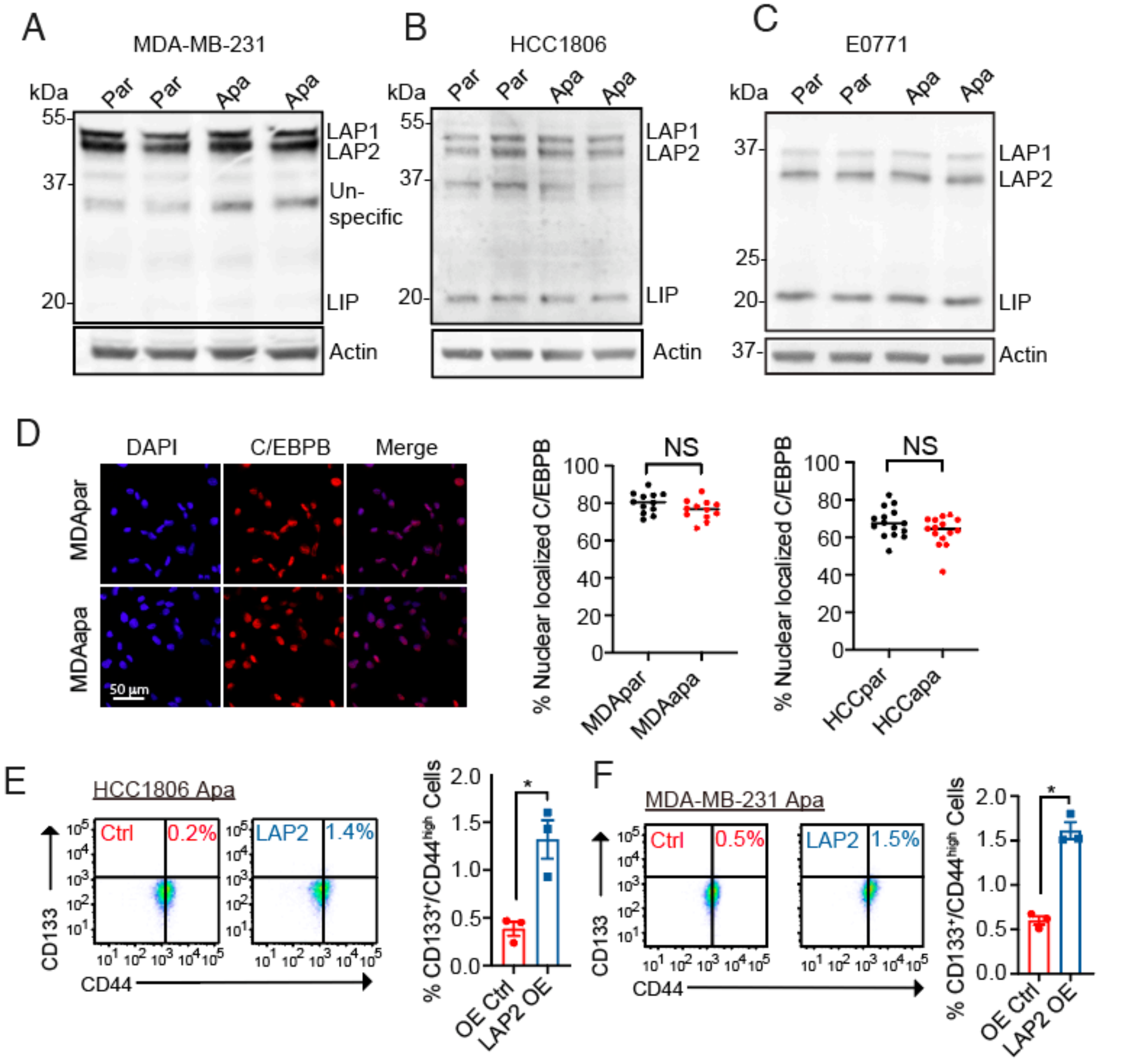


(A-C) Immunoblots of C/EBPB and Actin in parental and adapted MDA-MB-231 (A), HCC1806 (B) and E0771 (C) cell lines. Actin was used for the normalization.

(D) Representative images of C/EBPB-immunofluorescent staining on MDApar and MDAapa cells (left panel). Quantification (right panel) was calculated by the percentage of C/EBPB localized in the nucleus compared to the cytoplasm for MDA-MB-231 and HCC1806 parental and PA-adapted cell lines.

(E-F) $CD133^+/CD44^{high}$ cells population in adapted HCC1806 (E) and MDA-MB-231 (F) cells overexpressing LAP2. Cells were stained by CD133-APC and CD44-FITC antibodies and measured by flow cytometry. Quantification data is shown as mean ± SEM of three replicates.

For E-F, statistical significance determined with unpaired, two-tailed Student's t-test. (NS, P value > 0.05; *, P value < 0.05).

**Supplementary Figure 5.**

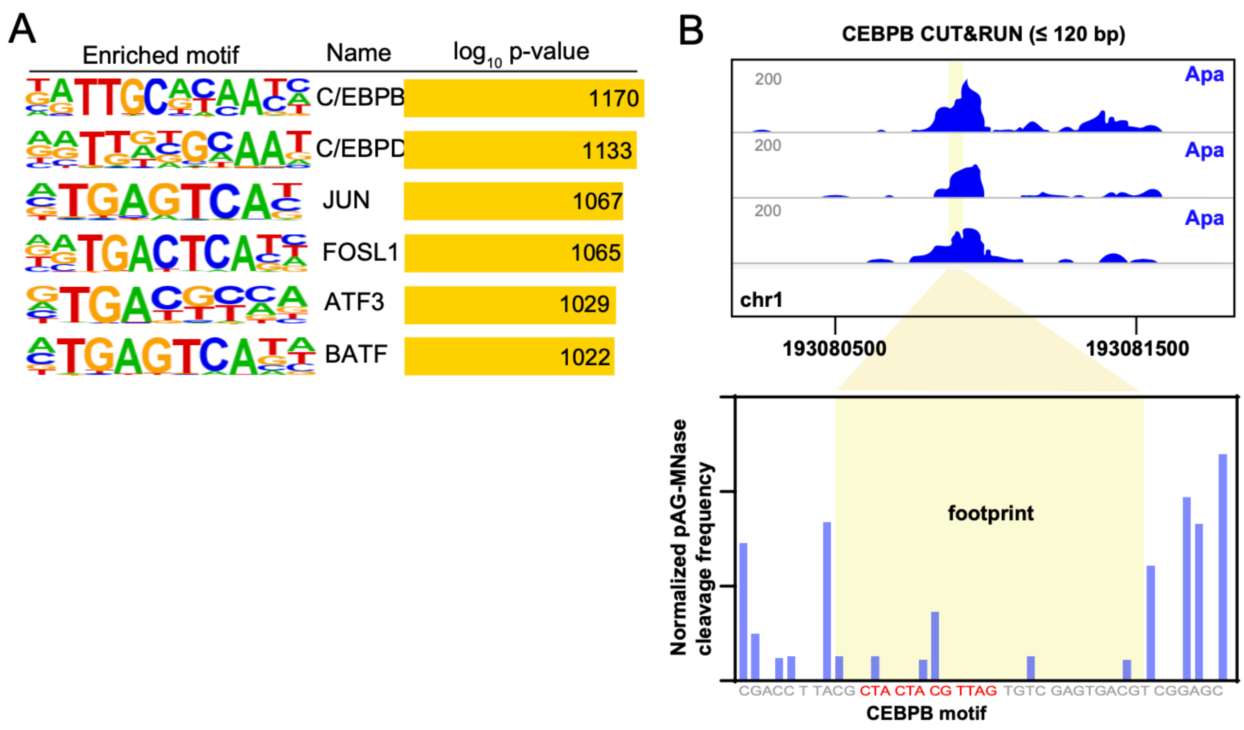


(A) Motifs enriched in C/EBPB Cut&Run footprints in MDAapa cells. The p-values shown in the figure were reported by HOMER using HOCOMOCO motifs.

(B) Single locus footprint analysis of C/EBPB Cut&Run experiments in adapted MDA-MB-231 cells. Upper panel shows representative genome browser tracks of C/EBPB Cut&Run signal in the specified region in chromosome 1 (chr1). Lower panel shows the total normalized pA/G-MNase cut frequency of the three biological replicates at each nucleotide around the *C/EBPB* motif within the identified footprint in the specified region.

**Supplementary Figure 6.**

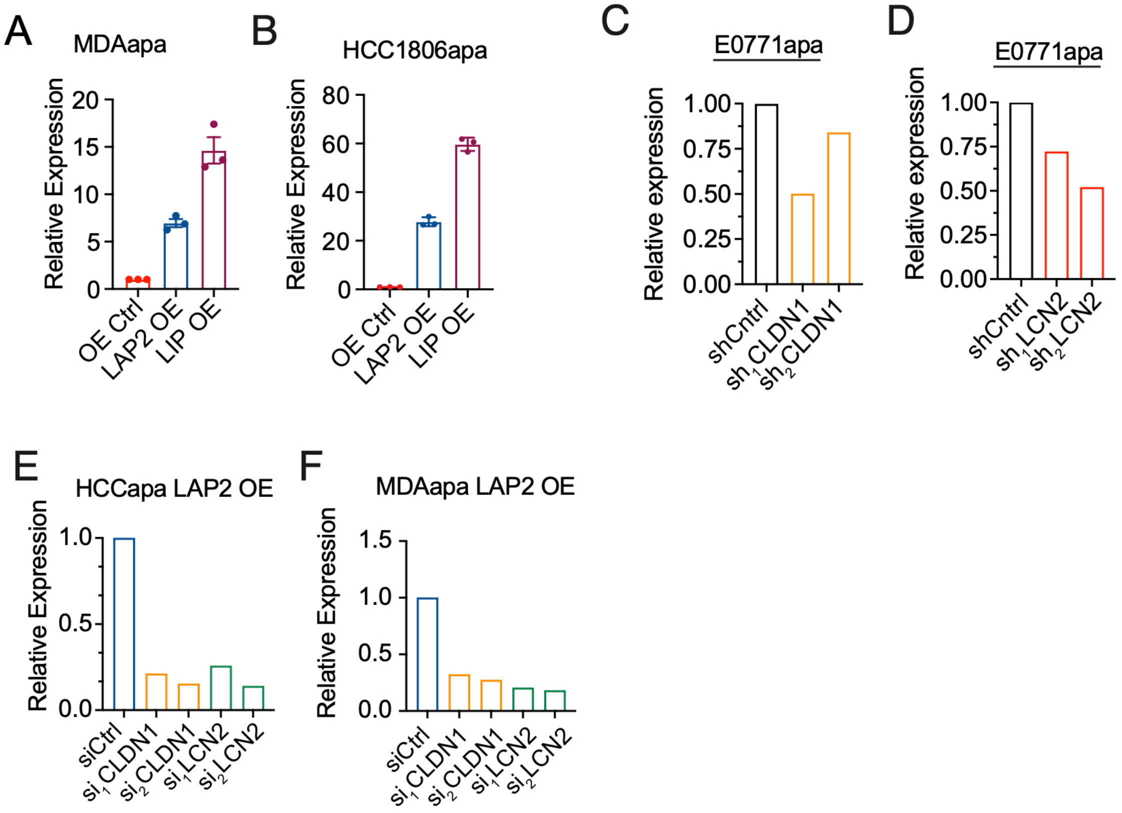


(A-B) RT-qPCR was used to measure changes in the expression of *C/EBPB* upon the overexpression of *C/EBPB LAP2* and *LIP* isoforms on adapted MDA-MB-231 (A) and HCC1806 (B) cells. The relative expression is shown as relative fold change over control cells. Data shown as mean ± SEM of three independently repeated experiments.

(C-D) RT-qPCR was used to measure efficiency of *Cldn1* (C) and *Lcn2* (D) knockdown in adapted E0771 cells. Knockdown was performed by using two independent shRNAs for each gene.

(E-F) RT-qPCR was used to measure efficiency of *CLDN1* (yellow) *and LCN2* (green) knockdown relative to knockdown control (siCtrl, blue) in adapted HCC1806 (E) and MDA-MB-231 (F) cells. Knockdown was performed by using two independent siRNAs for each gene.

**Supplementary Table S1**. List of genes included in the targeted sequencing of PM/ER$^-$/PR$^-$

| | | | |
|---|---|---|---|
| ABL1 | ERCC2 | MAPK10 | ROS1 |
| ABL2 | ERCC3 | MAPK7 | RPS6KB1 |
| ACVR2A | ERCC4 | MAPK8 | RPTOR |
| AKT1 | ERCC5 | MAPK9 | RRM2B |
| AKT2 | ESR1 | MCL1 | RSPO2 |
| AKT3 | ETV1 | MDM2 | RSPO3 |
| ALK | EZH2 | MDM4 | RUNX1 |
| APC | FADD | MED12 | SETD2 |
| AR | FAM123B | MED12L | SF3B1 |
| ARAF | FANCA | MED13 | SFTPA1 |
| ARFRP1 | FANCC | MED29 | SHC1 |
| ARID1A | FANCD2 | MEN1 | SKP2 |
| ARID1B | FANCE | MET | SLIT2 |
| ARID2 | FANCF | MITF | SMAD2 |
| ASXL1 | FANCG | MLH1 | SMAD3 |
| ATM | FAS | MLL | SMAD4 |
| ATR | FBXO11 | MLL2 | SMARCA4 |
| ATRX | FBXW7 | MLL3 | SMARCB1 |
| AURKA | FGFR1 | MPL | SMO |
| AURKB | FGFR2 | MRAS | SMURF1 |
| AXIN1 | FGFR3 | MRE11A | SOCS1 |
| BAG4 | FGFR4 | MSH2 | SOX10 |
| BAP1 | FH | MSH6 | SOX2 |
| BCL11A | FLT1 | MST1 | SOX9 |
| BCL2 | FLT3 | MTDH | SPOP |
| BCL2A1 | FLT4 | MTOR | SRC |
| BCL2L1 | FOXA1 | MUTYH | SRSF2 |
| BCL2L2 | FOXL2 | MYB | STAT3 |
| BCL6 | FOXO1 | MYC | STK11 |
| BCOR | FOXP4 | MYCL1 | SUFU |
| BIRC2 | GAB2 | MYCN | TBX22 |
| BIRC7 | GABRG1 | MYD88 | TBX3 |
| BLM | GATA1 | MYO3A | TERT |
| BPTF | GATA2 | MYO5B | TET2 |
| BRAF | GATA3 | MYOC | TGFBR2 |
| BRCA1 | GATA6 | NBN | TNFAIP3 |
| BRCA2 | GNA11 | NCOA2 | TOP1 |
| BRIP1 | GNAQ | NCOA3 | TP53 |
| BUB1B | GNAS | NF1 | TP63 |
| C11orf30 | GPC5 | NF2 | TP73 |
| CARD11 | GPR124 | NFE2L2 | TRAF2 |
| CASP8 | GRB2 | NGFR | TSC1 |
| CBL | GRB7 | NKX2-1 | TSC2 |
| CCND1 | GRID1 | NOTCH1 | TSHR |
| CCND2 | GUCY1A2 | NOTCH2 | U2AF1 |
| CCND3 | H3F3A | NOTCH3 | USP9X |
| CCNE1 | HIST1H3B | NOTCH4 | VEGFA |

**Supplementary Table S2**. Antibody panel used for mass cytometry analysis

| Isotope | Antigen | Cell Location | Epitope | Phenotype | Clone |
|---|---|---|---|---|---|
| 168Er | Axl | Extracellular | Total | Stemness/EMT | 1H12 |
| 160Gd | CD133 | Extracellular | Total, Epitope 1 | Stemness | AC133 |
| 173Yb | CD44 | Extracellular | Total, Surface | Stemness | IM7 |
| 158Gd | E-cadherin | Extracellular | CD324/E-Cadherin | Epithelial | "24E10" |
| 170Er | EGFR | Extracellular | Total EGFR | Epithelial organs/ initiates MAPk, Akt and JNk signalling | AY13 |
| 143Nd | N-cadherin | Extracellular | CD325/N-Cadherin | Stemness/mesenchymal | 8C11 |
| 156Gd | p38 | Intracellular | p38 [T180/Y182] | MAPK for stress response | D3F9 |
| 152Sm | pAkt | Intracellular | pAkt [S473] | PI3K pathway | D9E |
| 176Yb | pCreb | Intracellular | pCREB [S133] | Transcription factor- stress and growth | 87G3 |
| 151Eu | pEGFR | Intracellular | pEGFR [Y1068] | Activated EGFR | Y38 |
| 154Sm | pErk1/2 | Intracellular | pT202/pY204 | Branch of MAPK-Mek pathway | 20A |
| 175Lu | pHistone H3 | Intracellular | pHistone H2A.X [Ser139] | Metaphase. Activated dwnstream of p38 or Erk1/2 | HTA28 |
| 159Tb | pMAPKAPK2 | Intracellular | pMAPKAPK2 [T334] | ERk1/2 activated protein downstream of p38. Response to stress | 27B7 |
| 166Er | pNFKB | Intracellular | pNF-kB p65 [S529] | Transcription Factor mediator of inflammatory and immune responses | K10-895.12.50 |
| 162Dy | pPLCgamma 2 | Intracellular | pPLCg2 [pY759] | Mediator of inflammatory and immune responses | K86-689.37 |
| 150Nd | pRb | Intracellular | pRb [S807/811] | G1 to S cell cycle phase | J112-906 |
| 172Yb | pS6 | Intracellular | pS6 [S235/S236] | Protein translation | N7-548 |
| 141Pr | pSHP2 | Intracellular | Y580 | RTK phosphatase promotes signaling of JAK/STAT, PI3K/Akt Ras/MAPK pathway | D66F10 |
| 153Eu | pStat1 | Intracellular | Y704 | | 4a |
| 145Nd | pStat3 | Intracellular | pY705 | | 4/p |
| 146Nd | pStat5 | Intracellular | pY694 | | 00047 |
| 149Sm | pStat6 | Intracellular | Y641 | | 18/P-stat6 |
| 163Dy | TGFbeta | Intracellular | Total | | TW4-6H10 |
| 154Sm | Vimentin | Intracellular | Total | Mesenchymal | D21H3 |
| 167Er | YAP | Intracellular | CTD 379-407 | Stemness Hippo | H9 |
| 172YT | CC3 | Intracellular | Cleavage at D175 | Apoptosis | 5A1E |
| 164DY | CK7 | Intracellular | Total | Luminal marker | RCK105 |

**Supplementary Table S3**. PCR Primer sequences used for barcode amplification

| | Sequence | Length |
|---|---|---|
| **WS PCR Forward Primer** | AATGATACGGCGACCACCGAGATCTACACACTGACTGCAGTCTGAGTCTGACAG | 54 |
| **WS_Rev_Index_011** | CAAGCAGAAGACGGCATACGAGATGTATCACGACGTGACTGGAGTTCAGACGTGTGCTCTTCCGATCTCTAGCACTAGCATAGAGTGCGTAGCT | 94 |
| **WS_Rev_Index_013** | CAAGCAGAAGACGGCATACGAGATAGCGTCTGATGTGACTGGAGTTCAGACGTGTGCTCTTCCGATCTCTAGCACTAGCATAGAGTGCGTAGCT | 94 |
| **WS_Rev_Index_014** | CAAGCAGAAGACGGCATACGAGATCAGCATGTCTGTGACTGGAGTTCAGACGTGTGCTCTTCCGATCTCTAGCACTAGCATAGAGTGCGTAGCT | 94 |
| **WS_Rev_Index_015** | CAAGCAGAAGACGGCATACGAGATGTACTCATCGGTGACTGGAGTTCAGACGTGTGCTCTTCCGATCTCTAGCACTAGCATAGAGTGCGTAGCT | 94 |
| **WS_Rev_Index_016** | CAAGCAGAAGACGGCATACGAGATTCTGCAGCTAGTGACTGGAGTTCAGACGTGTGCTCTTCCGATCTCTAGCACTAGCATAGAGTGCGTAGCT | 94 |
| **WS_Rev_Index_017** | CAAGCAGAAGACGGCATACGAGATACTGTACTCGGTGACTGGAGTTCAGACGTGTGCTCTTCCGATCTCTAGCACTAGCATAGAGTGCGTAGCT | 94 |
| **WS_Rev_Index_018** | CAAGCAGAAGACGGCATACGAGATCGACAGCTATGTGACTGGAGTTCAGACGTGTGCTCTTCCGATCTCTAGCACTAGCATAGAGTGCGTAGCT | 94 |
